# PARP1 catalytic domain mutations drive high-level resistance to saruparib while preserving DNA damage response vulnerabilities

**DOI:** 10.64898/2026.05.08.723870

**Authors:** Matthew R. Jordan, Jessica L. Kersey, Joy E. Garrett, Sheng Liu, Jun Wan, John J. Turchi

## Abstract

Clinical poly (ADP)-ribose polymerase (PARP) inhibitors (PARPi) are limited by toxicities associated with inhibition of multiple PARP family proteins and acquired resistance. As PARP1-specific inhibitors, like saruparib (AZD5305), move toward standard-of-care status for BRCA and HR-deficient cancers replacing less specific PARPi, defining mechanisms of intrinsic and acquired resistance is essential for developing effective treatment strategies. Here, we established 5 saruparib-resistant (SR) cell lines from BRCA1-deficient MDA-MB-436 triple negative breast cancer (TNBC) cells using a selection strategy of high-level dosing consistent with clinical exposure, yielding models that are >1,000-fold resistant to saruparib. Whole genome sequencing identified PARP1 catalytic domain mutations in all SR cell lines, and *in vitro* reconstitution of these PARP1 mutants confirmed them as drivers of saruparib resistance, in contrast to HR restoration as observed in the case of less-selective PARPi. PARP1 mutations also induce altered saruparib-dependent PARP1 trapping and PARylation inhibition. While these mutations render cells highly resistant to saruparib, differential sensitivity to other PARPi was observed and SR cell lines retain, and in some cases, increase, sensitivity to alternative clinical PARPi and DNA damage response (DDR)-targeted therapeutics. Our findings demonstrate that high-intensity selection pressure favors target-site mutation over pathway restoration as a primary escape mechanism from PARP1-selective inhibition. This study provides a first-in-class characterization of saruparib resistance and maps a clear therapeutic path forward. By identifying these specific PARP1 mutations and their collateral DDR vulnerabilities, we provide the molecular framework necessary to monitor and treat patients who progress on next-generation PARP1-selective inhibitors.

## Introduction

Loss of function mutations in breast cancer associated genes 1 and 2 (BRCA1 and BRCA2) lead to a significant predisposition to malignancy in numerous cancers, notably for their namesake breast cancer [1]. As both BRCA1 and BRCA2 are essential for the high-fidelity double-strand DNA break (DSB) repair pathway homologous recombination (HR), this predisposition is attributed to the accumulation of additional, oncogenic mutations in the absence of functioning HR and the reliance on more error prone DSB repair pathways. Despite these factors enabling tumor progression, BRCA deficiency also provides a vulnerability that was exploited by the use of poly (ADP)-ribose polymerase (PARP) inhibitors (PARPi) that cause synthetic lethal killing of BRCA-deficient cancers [2,3]. In 2014, olaparib became the first FDA approved PARPi for the treatment of BRCA-deficient cancers, and targeted PARPi therapy was eventually expanded to other HR-deficient contexts. There are currently four FDA-approved PARPi used for the treatment of BRCA-deficient cancers in the clinic: olaparib, niraparib, rucaparib, talazoparib [4].

The PARP family of proteins consists of 17 members that function in a myriad of pathways [5]. PARP1 performs ∼90% of PARP function in the DNA damage response (DDR), with PARP2 responsible for the majority of the remaining ∼10% [6,7]. PARP1 participates in numerous DNA repair pathways and is important for the repair of DNA single-stranded breaks (SSBs) as the major cellular SSB sensor. PARP1 contains four DNA binding domains, one BCRT domain, and a catalytic domain comprised of a helical domain (HD) and the ADP-ribosyltransferase (ART) domain [8]. In the absence of DNA, the HD is autoinhibitory and precludes substrate NAD^+^ binding in the ART domain [9]. Upon DNA binding, the HD becomes unfolded via allosteric linkage with the DNA binding domains, which activates its enzymatic activity to poly ADP-ribosylate (PARylate) itself, surrounding proteins, and DNA [8,9]. PARP1 auto-PARylation then releases PARP1 from the SSB, and other DNA repair proteins that possess PAR-binding motifs are simultaneously recruited to the site of damage to repair the DNA lesion [10]. More than two decades after the discovery, the toxic event induced by PARPi in BRCA-deficient cells remains an open question. The classical model proposes that upon PARP inhibition, unrepairable SSBs are converted to one-ended DSBs during DNA replication. In the absence of a functioning HR pathway, repair of the one-ended DSB either cannot occur or must go through more error prone pathways, which is ultimately detrimental to genomic stability and leads to cell death [11,12]. However, BRCA proteins have also been shown to function in replication fork protection and ssDNA gap suppression, where PARPi treatment ultimately results in DSB formation from the degradation of reversed replication forks and the conversion of ssDNA gaps into DSBs after replication run-through, respectively [13–16]. In addition, PARPi block PARP1 auto-PARylation which therefore traps PARP1 on the DNA lesion rather than being released upon enzymatic activation, potentially becoming an impediment [17]. Therefore, the question as to whether catalytic inhibition of PARP proteins or PARP trapping is causing toxicity is also debated in the field [18–21]. Despite the clinical success of PARPi in treating BRCA-deficient and other HR-deficient cancers, around 30-40% of BRCA-deficient patients are intrinsically resistant and do not initially respond to PARPi [22]. Of those that do initially respond, 50-70% ultimately develop PARPi resistance [22]. PARPi resistance is thus a major clinical burden, and understanding the exact mechanisms driving PARPi resistance is essential for designing rationally targeted therapeutic strategies to overcome resistance. Known mechanisms of acquired resistance can generally be described as restoring/compensating for BRCA function, affecting drug metabolism, or altering PARP1 stability or ADP-ribosylation. Most commonly observed in the clinic (≥75%), HR can be restored in BRCA-deficient cells through a few distinct mechanisms: BRCA reversion mutations can re-establish the BRCA reading frame and restore BRCA function, loss of DSB end-binding factors can restore HR end resection independent of BRCA expression, and upregulation of other HR factors can compensate for the loss of BRCA and restore HR activity [16,23–25]. Additionally, PARPi efflux via upregulation of a variety of transporters confers PARPi resistance [26]. While rare, a PARP1 mutation that alters the DNA sensing allosteric activation of ADP-ribosylation has been observed to result in PARPi resistance [27]. Moreover, the loss of the PAR eraser poly ADP-ribose glycohydrolase (PARG) also promotes PARPi resistance [28].

Current clinical PARPi are nonspecific to the PARP family of proteins, which contributes to clinical side effects and toxicity that has been attributed to PARP2 inhibition in healthy tissues [29,30]. Saruparib (AZD5305) is a recently developed PARPi that is selective for PARP1 and therefore exhibits significantly improved tolerability, is anticipated to be the standard of care PARPi, and is currently undergoing phase 3 clinical trials [31,32]. Additionally, saruparib treatment has proven efficacious in systems of acquired resistance to other clinical PARPi when combined with ATR inhibitors (ATRi) or platinum-based therapies [32–34].

In this study, we characterized resistance mechanisms to the PARP1-specific inhibitor saruparib by generating five BRCA1-deficient triple-negative breast cancer (TNBC) cell lines with acquired saruparib resistance. Across these cell lines, we identified three PARP1 mutations that confer >1,000-fold resistance to saruparib, but do not exhibit pan-PARPi resistance. Reconstitution of these mutations *in vitro* recapitulates the differences in cellular viability suggesting that these PARP1 mutations are driving saruparib resistance. Saruparib-resistant cell lines exhibit altered PARylation dynamics with and without exogenous DNA damage and altered PARP trapping. While it is unclear whether saruparib resistance arises from the loss of MAR/PARylation or PARP1 trapping, changes in talazoparib sensitivity appear to arise from alterations in PARP1 trapping and not MAR/PARylation. Lastly, we find that saruparib-resistant cells remain sensitive to other common breast cancer chemotherapeutics and DNA damage response-targeted agents. These results suggest PARP1 mutations are drivers of resistance to PARP1-specific inhibitors and demonstrate potential therapeutic strategies for patients who progress on saruparib.

## Results

### Generation of saruparib-resistant BRCA1-deficient TNBC cell lines

As saruparib is anticipated to be the standard of care PARPi for patients with BRCA/HR-deficient tumors, we aimed to generate saruparib-resistant cell lines in order to characterize resistance mechanisms that may present in future patients. Additionally, we hypothesized that resistance mechanisms derived from the PARP1-specific inhibitor may well be different from known mechanisms of PARPi resistance in systems of pan-PARP inhibition. To this end, we used the BRCA1-deficient MDA-MB-436 TNBC cell line model. To mimic the clinical setting, we chose to treat MDA-MB-436 cells with a relatively high dose ∼20 times greater than the half-maximal inhibitory concentration (IC50), rather than maintaining cultures with a lower, slow dose escalation that has been used in other PARPi resistance models [23,34–36] (Figure 1A & S1A). Importantly, this dose (30 nM saruparib), is insufficient to generate off-target inhibition of PARP2, as can occur at concentrations exceeding 0.5 µM [32]. Moreover, this concentration is comparable to the murine plasma concentration resulting from a saruparib dosage that induced >90% tumor regression in mouse models and is therefore representative of the clinical setting [32]. After ∼5 weeks, we had selected for 10 saruparib-resistant colonies, of which 5 were successfully expanded into stable cell lines. These cell lines are henceforth referred to as 436-SR-1, 436-SR-4, 436-SR-5, 436-SR-9, and 436-SR-10 for MDA-MB-<u>436</u> <u>S</u>aruparib-<u>R</u>esistant (SR) cell lines.

**Figure 1.**
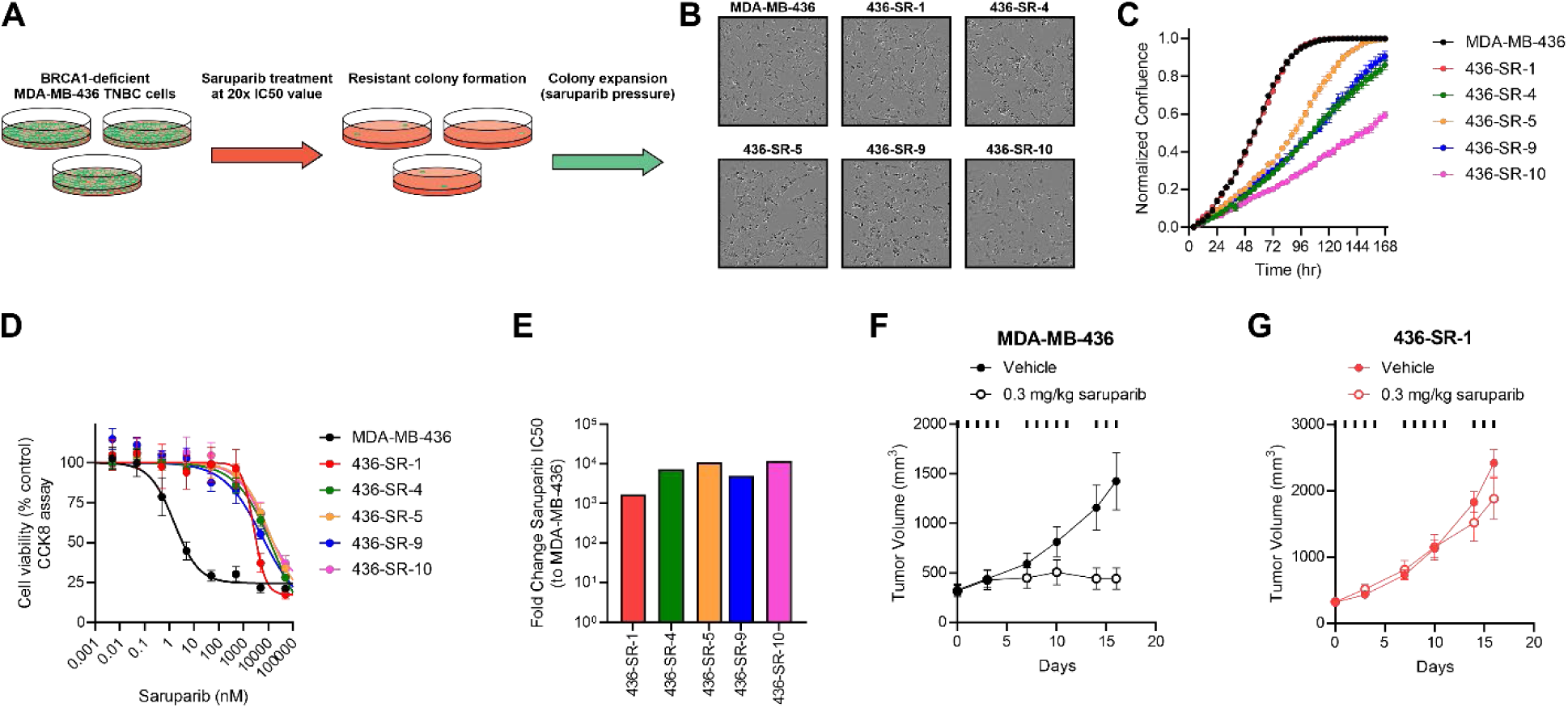
Generation of saruparib-resistant BRCA1-deficinet TNBC cell lines. **A)** Experimental scheme for the selection of saruparib-resistant cell lines derived from MDA-MB-436 cells. **B)** Comparison of cell morphology of MDA-MB-436 and SR cell lines. **C)** Normalized confluence over time for MDA-MB-436 and SR cells as measured by Incucyte live-cell imaging. **D)** Dose-response curves for MDA-MB-436 and SR cells treated with saruparib. **E)** SR cell lines’ fold-changes in saruparib IC50 values relative to parental MDA-MB-436 cells derived from the data in panel D. **F)** Assessment of tumor growth of MDA-MB-436 cells and **G)** 436-SR-1 cells subcutaneously implanted in NSG mice. Mice were treated via oral gavage with doses indicated by the solid bars. Data in panels C, D, F, and G are depicted as mean ± SEM.

All SR cell lines have largely similar cellular morphology as compared to parental MDA-MB-436 cells, with the exception of subpopulations of 436-SR-9 and 436-SR-10 cells that have larger, rounded morphology rather than spindly (Figure 1B). 436-SR-1 cell growth kinetics are essentially identical to parental MDA-MB-436 cells, while the remaining 4 cell lines exhibit a slower growth phenotype as measured via live cell imaging (Figure 1C). Importantly, whereas saruparib exhibits an IC50 value of 1.6 nM for parental MDA-MB-436 cells, all SR cell lines exhibit >1,000-fold resistance to saruparib (Figure 1D-E), at which off-target effects are likely responsible for cell death [32]. To confirm resistance in a tumor model, we implanted NOD-scid/IL2Rg-null (NSG) mice with either MDA-MB-436 cells or 436-SR-1 cells as a representative of SR cell lines as they exhibited similar growth kinetics (Figure 1C). Tumors were allowed to grow to a relatively large size of ∼325 mm^3^ and were randomized to vehicle or 0.3 mg/kg saruparib treatment regimens 5 days on/2 days off by oral gavage. Tumor volume and terminal tumor weight measurements demonstrate flat-lining of MDA-MB-436 tumor growth, while 436-SR-1 tumors indeed remain resistant to saruparib treatment (Figure 1F-G & S1B), validating saruparib resistance *in vivo*.

### SR cell lines do not exhibit pan-PARPi resistance or HR restoration

With the establishment of *in vitro* and *in vivo* resistance to the PARP1-specific inhibitor saruparib, we next examined the sensitivity of SR cell lines to other pan-PARPi as compared to parental MDA-MB-436 cells; we utilized 3 clinical PARPi (olaparib, talazoparib, and niraparib) and 1 pre-clinical PARPi that also inhibits tankyrase 1 & 2 (stenoparib). Importantly, other PARPi-resistant cell lines in the literature, typically selected for through slow dose escalation, exhibit broad range PARPi resistance [23,34–36], as would be expected for common resistance mechanisms of direct BRCA1 reversion or Shieldin/53BP1 loss that would restore HR. Intriguingly, we found that this was not the case for SR cell lines, with some cell lines actually more sensitive to other PARPi (Figure 2A-E). While the 5 SR cell lines can be grouped into those that are generally more sensitive to other PARPi (436-SR-1, 436-SR-4, 436-SR-5) and those that are generally more resistant (436-SR-9, 436-SR-10), four of the five SR cell lines exhibit differential sensitivity to the four PARPi tested and only 436-SR-10 cells exhibit a broad range response (Figure 2E). Sensitivity or resistance was on the order of 10-fold to 100-fold change in IC50 (Figure 2E), significantly less than what was observed with saruparib (Figure 1E). These differential sensitivity results suggest a resistance mechanism unrelated to the restoration of HR. Indeed, like parental MDA-MB-436 cells, all 5 SR cell lines do not possess HR activity as compared to HR-competent, BRCA1-wild-type MDA-MB-468 cells in an extrachromosomal reporter assay (Figure 2F).

**Figure 2.**
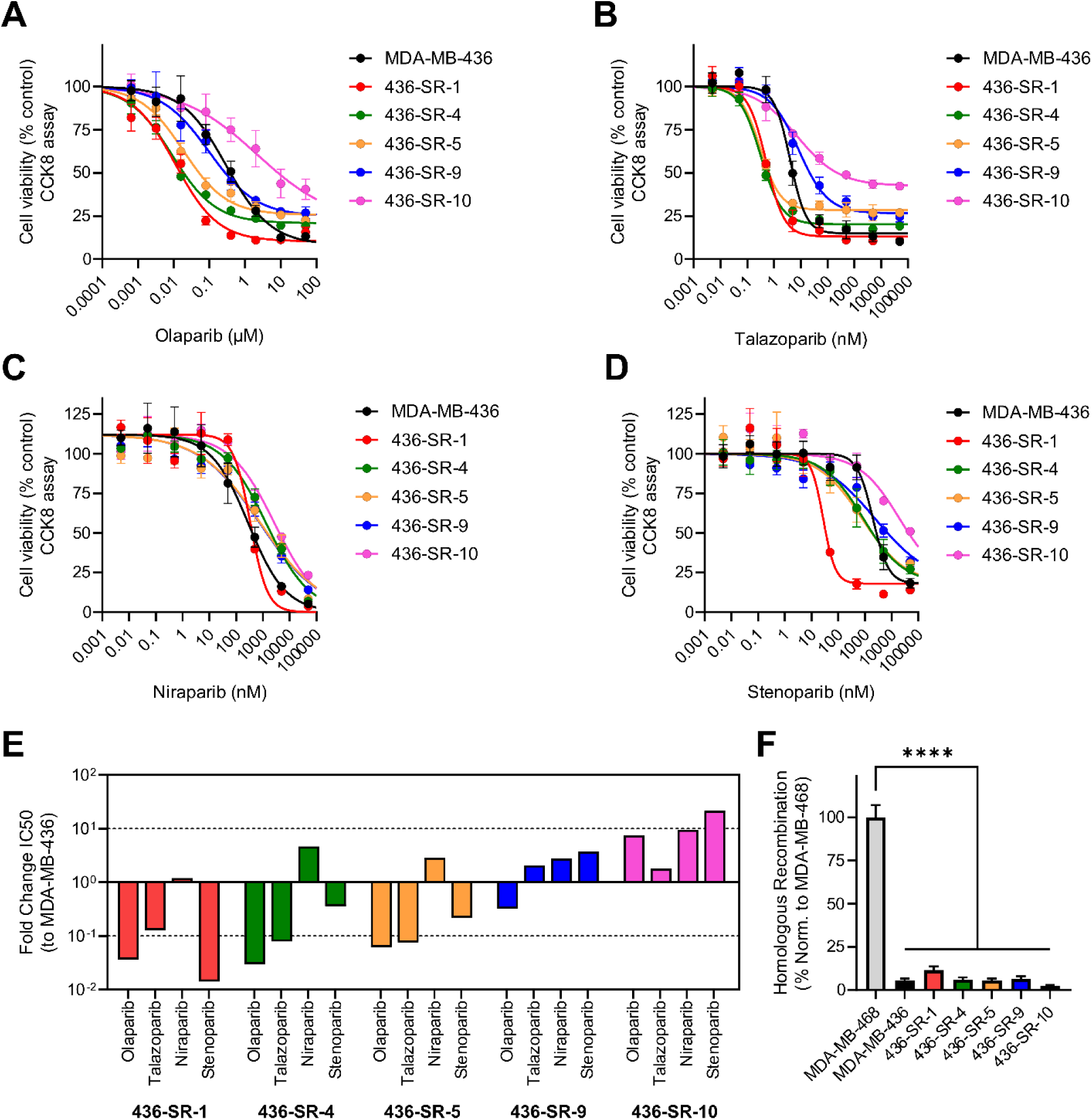
SR cell lines do not exhibit pan-PARPi resistance or HR restoration. Dose-response curves for MDA-MB-436 and SR cells treated with **A)** olaparib, **B)** talazoparib, **C)** niraparib, or **D)** stenoparib. **E)** SR cell lines’ fold-changes in olaparib, talazoparib, niraparib, and stenoparib IC50 values relative to parental MDA-MB-436 cells derived from the data in panels A-D. **F)** Measurement of homologous recombination activity using an extrachromosomal reporter assay. BRCA1-proficient MDA-MB-468 cells were used as a positive control. Data in panels A-D and F are depicted as mean ± SEM from triplicate experiments. **** P < 0.0001.

### All SR cell lines possess PARP1 catalytic domain mutations

To investigate the mechanisms of resistance for each cell line, we performed whole genome sequencing (WGS) and RNA-seq analysis compared to the parental MDA-MB-436 cell line (Table S1 & S2). In both data sets, we found strong overlap between 436-SR-4 and 436-SR-5 cell lines, and 436-SR-9 and 436-SR-10 cell lines, suggesting a unified, but divergent ancestry between each pair of cell lines (Figure S2A-B). We found no evidence of common PARPi resistance mechanisms such as BRCA1 reversion, loss of 53BP1/shieldin complex, or loss of PARG expression (Table S2). In addition, pathway analysis of the RNA-seq data for each cell line did not reveal any notable gene expression alterations known or predicted to be involved in PARPi resistance (Figure S2C). Strikingly, all five SR cell lines contained heterozygous mutations in the PARP1 catalytic domain: G871W in 436-SR-1, K893E in 436-SR-4 and 436-SR-5, and D770H in 436-SR-9 and 436-SR-10 (Figure 3A-C & S2D-E). D770 is located within the auto-inhibitory HD and is part of an acidic triad that points into the active site to preclude NAD^+^ binding in the absence of DNA [9] (Figure 3B-D). G871 is within an α-helix in the ART domain in close proximity to D770 (Figure 3B-D). The G871W mutation likely results in protrusion of the bulky tryptophan side chain into the NAD^+^ binding site, as modeled by AlphaFold3, that may impact substrate binding (Figure 3D). Moreover, some PARPi, like saruparib and olaparib (Figure 3C-D, gray and orange sticks, respectively), that extend into this HD-ART interface may exhibit perturbed binding to PARP1 and thus lessened efficacy, as opposed to those PARPi that primarily bind within the base of the PARP1 NAD^+^ binding site, like talazoparib and niraparib (Figure 3C-D, magenta and yellow sticks, respectively). K893E is located in the ART domain and forms three hydrogen bonds to the amide backbone of a loop that forms the HD-ART interface at the top of the NAD^+^ binding site (Figure 3B,E). The loss of stabilization of this loop by the charge swapping K893E mutation may impact the auto-inhibitory function of the HD.

**Figure 3.**
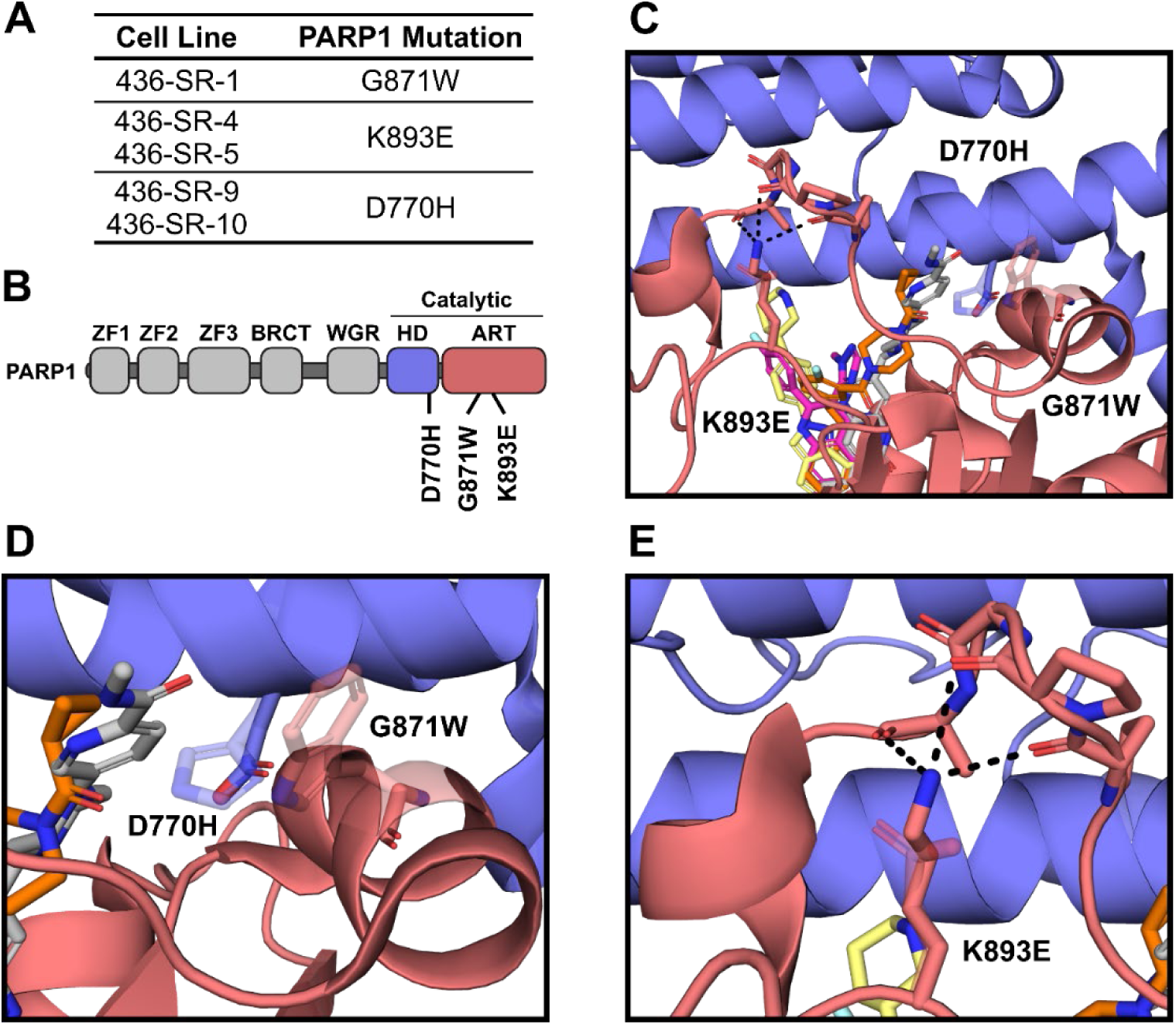
All SR cell lines possess PARP1 catalytic domain mutations. A) PARP1 catalytic domain mutations identified from WGS (see Table S1). B) Domain structure of PARP1. Throughout the figure, the helical domain (HD) is colored blue and the ADP-ribosyl transferase (ART) domain is colored red. C) Structure of the NAD^+^ binding site within the active site of the catalytic domain of PARP1 bound to saruparib (gray, PDB: 9ETQ), olaparib (orange, PDB: 7AAD), talazoparib (magenta, PDB: 7KK3), and niraparib (yellow, PDB: 7KK5) generated by aligning catalytic domains from each referenced structure to the saruparib-bound structure. AlphaFold3 models for each PARP1 mutation identified in this study were also aligned with each individual WT residue and mutation displayed as sticks. D) Region of D770H and G871W mutations at the interface of the HD and ART. E) Region of K893E mutation that forms hydrogen bonds (black dashes) with a flexible loop that forms the interface with the HD. In panels C-E, WT residues are solid, while mutated residues are semitransparent.

### In vitro characterization of SR PARP1 mutations recapitulates cellular sensitivity

The occurrence of these three catalytic domain mutations from the five generated SR cell lines led us to hypothesize that they were the defining genetic alteration driving saruparib resistance. It is notable that all three mutations are located around the periphery of the NAD^+^/PARPi binding pocket. Though these residues do not make any direct contacts with saruparib, the mutations may alter the general structure of the binding site and impact the ability of saruparib to inhibit PARP1. To address this possibility, we utilized an *in vitro* assay using the PARP1 661-1014 L713F construct, a constitutively active PARP1 catalytic domain henceforth referred to as PARP1-CAT [37]. We purified PARP1-CAT, as well as the three identified mutations, and assessed enzyme activity by measuring auto-PARylation activity via Western blotting. K893E PARP1-CAT exhibited comparable baseline levels of auto-PARylation, however both G871W PARP1-CAT and particularly D770H PARP1-CAT exhibited reduced baseline levels of auto-PARylation (Figure S3A-B). We next assessed the inhibitory potency of saruparib for each PARP1-CAT protein through the same experimental procedures and found that all three mutant proteins exhibited significant saruparib resistance (Figure 4A-B). G871W PARP1-CAT enzyme activity was most resistant to saruparib with a 50-fold greater IC50 value, followed by D770H PARP1-CAT and K893E PARP1-CAT that were 13-fold and 7-fold more resistant to saruparib inhibition than PARP1-CAT, respectively (Figure 4A-B). Interestingly, while the three mutant proteins exhibited resistance to saruparib inhibition of auto-PARylation, resistance was significantly less than what was observed in cell viability assays of the SR cell lines with heterozygous expression of the mutant proteins (Figure 1D-E). We also assessed the *in vitro* potency of talazoparib for each protein and observed differential sensitivity, though to a smaller degree than inhibition by saruparib (Figure 4C-D). G871W PARP1-CAT was 6-fold more sensitive to talazoparib inhibition, while D770H PARP1-CAT was inhibited similarly to PARP1-CAT, and K893E PARP1-CAT was ∼2-fold more resistant to talazoparib (Figure 4C-D).

**Figure 4.**
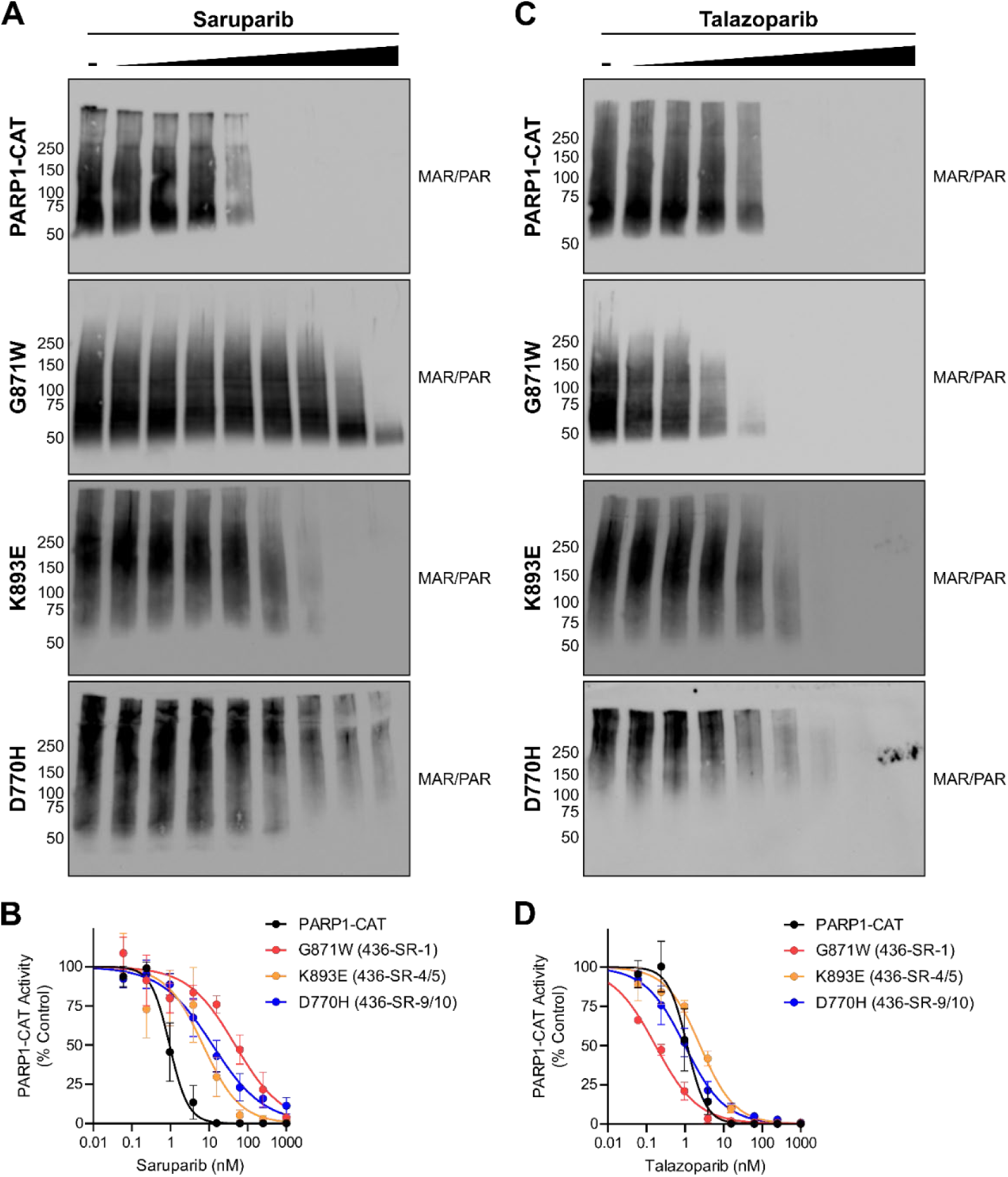
In vitro characterization of SR PARP1 mutations recapitulates cellular sensitivity. **A)** Representative Western blots of auto-MAR/PARylation activity of purified PARP1-CAT and SR mutants treated with DMSO control or increasing concentrations of saruparib. **B)** Quantification of the inhibition of PARP1-CAT activity depicted in panel A from triplicate measurements. **C)** Representative Western blots of auto-MAR/PARylation activity of purified PARP1-CAT and SR mutants treated with DMSO control or increasing concentrations of talazoparib. **D)** Quantification of the inhibition of PARP1-CAT activity depicted in panel C from triplicate measurements. Data in panels B and D are depicted as mean ± SEM from triplicate experiments.

### SR cell lines exhibit altered MAR/PARylation and PARP1 trapping in response to PARPi

We next assessed PARP1 and PARP2 expression in the SR cell lines, as well as the PARP-directed response to DNA damage. Cells were treated with or without 0.1 mg/mL methyl methanesulfonate (MMS), a potent methylating agent that induces SSBs and PARP activation, for 2 hours before soluble protein fractions were visualized by Western blotting (Figure 5A). We observed somewhat reduced levels of PARP1 expression in all SR cell lines, and minor fluctuations in PARP2 expression (Figure 5A & S4A-B). Upon DNA damage induction, PARP1 recruitment occurs rapidly (∼5-10 seconds), followed by massive PARylation at the damage site [38,39]. PARP1 removal from the site of damage occurs after ∼5 minutes [40] as other repair factors are recruited and the lesion is repaired. By ∼15 minutes post-DNA damage, the majority of PARylation has been removed primarily by PARG [38]. Consistently, under our experimental conditions, the MAR/PARylation levels of MDA-MB-436 cells are greatly reduced 15 minutes after MMS treatment and gradually increase over 2 hours, though are not completely restored to initial levels (Figure S4C). Therefore, any changes in MAR/PARylation after 2 hours of MMS treatment are indicative of a PARP-dependent response to damaged DNA. There are two striking observations in the comparison of MAR/PARylation levels across SR cell lines 2 hours post-MMS treatment. First, 436-SR-1 cells have significantly elevated MAR/PARylation despite reduced PARP1 expression, both basally and in response to DNA damage (Figure 5A & S4D), despite the G871W mutation resulting in reduced activity *in vitro* (Figure S3). Second, 436-SR-9 and 436-SR-10 cells exhibit no response to DNA damage, as MMS does not cause any change in MAR/PARylation, and may indicate that the D770H mutation results in constitutive PARP activity that is potentially uncoupled from DNA damage sensing (Figure 5A & S4D).

**Figure 5.**
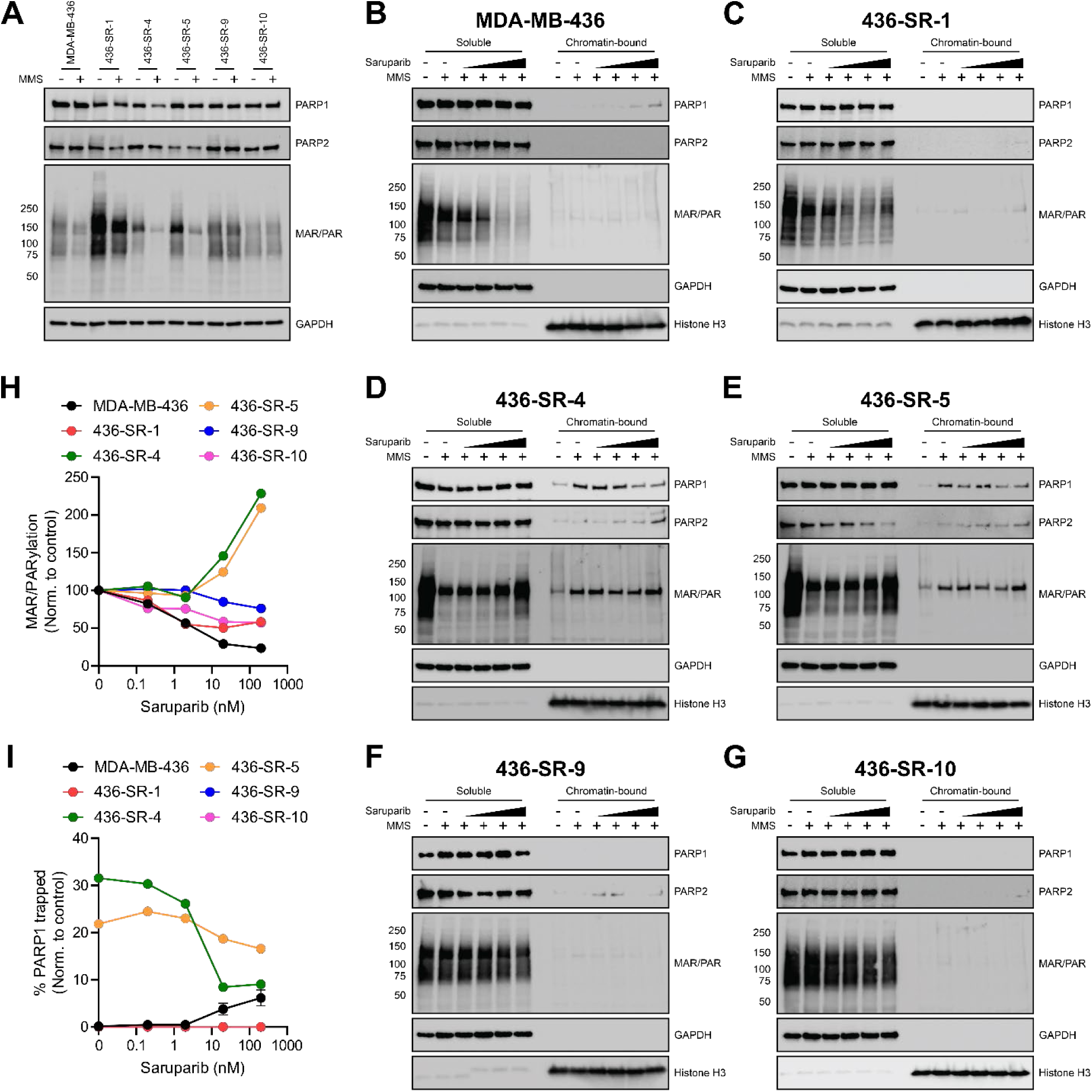
SR cell lines exhibit altered MAR/PARylation and PARP1 trapping in response to saruparib. **A)** Representative Western blots of PARP1 and PARP2 expression and MAR/PARylation from the soluble fraction of MDA-MB-436 and SR cells in response to 0.1 mg/mL MMS treatment for 2 hours. Representative Western blots of soluble and chromatin-bound fractions from **B)** MDA-MB-436, **C)** 436-SR-1, **D)** 436-SR-4, **E)** 436-SR-5, **F)** 436-SR-9, and **G)** 436-SR-10 cells treated with DMSO control, 0.1 mg/mL MMS, and/or 0.2, 2, 20, 200 nM saruparib for 2 hours. **H)** Quantification of saruparib inhibition of MAR/PARylation in MDA-MB-436 and SR cells from panels B-G. **I)** Quantification of saruparib-induced PARP1 trapping in MDA-MB-436 and SR cells from panels B-G.

There is much debate in the field as to whether PARPi toxicity arises from inhibition of PARP catalytic activity or PARP trapping [18–21], and thus saruparib resistance could potentially arise from the loss of either mechanism. We therefore assessed the levels of MAR/PARylation and PARP1 trapping in response to saruparib in all SR cell lines compared with the parental MDA-MB-436 cell line. Cells were treated with MMS to induce DNA damage to be sensed by PARP1 and were co-treated with increasing amounts of saruparib for 2 hours before separation of soluble and chromatin bound fractions for western blot analysis as indicated by GAPDH or Histone H3, respectively (Figure 5B-G). As expected, control MDA-MB-436 cells exhibit dose-dependent inhibition of MAR/PARylation activity and observable trapping of a small fraction of PARP1 at higher concentrations of saruparib (Figure 5B). Interestingly, 436-SR-1 cells similarly exhibited inhibition of MAR/PARylation, though with some distinctions (Figure 5C). Inhibition of MAR/PARylation in 436-SR-1 cells is maximal at 2 nM saruparib, with no further decrease up to 200 nM saruparib, however this residual MAR/PARylation at elevated saruparib concentrations is higher than in MDA-MB-436 cells (Figure 5C,H). More strikingly, however, 436-SR-1 cell lines do not exhibit any PARP1 trapping (Figure 5C). Taken together, the G871W mutation abolishes saruparib-dependent PARP1 trapping and its enzymatic activity is sustained under saruparib treatment, though modestly inhibited.

Both 436-SR-4 and 436-SR-5 cells exhibited similar, unexpected responses to saruparib treatment (Figure 5D-E). After the initial reduction in MAR/PARylation in the response to MMS-induced DNA damage, MAR/PARylation levels strikingly increased with increasing saruparib treatment (Figure 5D-E, H). Moreover, PARP1 trapping was very high in cells treated with MMS in the absence of saruparib, and saruparib treatment resulted in a dose-dependent decrease in PARP1 chromatin retention (Figure 5D-E, I). Congruous with this elevated level of MMS-induced PARP1 trapping in the absence of any PARPi, a persistent MAR/PARylated band that is consistent with the molecular weight of auto-PARylated PARP1 is retained in the chromatin-bound fractions under all MMS-treated conditions, a phenomenon not observed in any of the other examined cell lines (Figure 5D-E). This suggests some sort of impairment in the pro-release function of auto-PARylation which ultimately leads to significant PARP1 trapping upon DNA damage events. Importantly, despite the decrease in trapped PARP1 upon titration with saruparib relative to the control, 436-SR-4 and 436-SR-5 cells actually display higher levels of trapped PARP1 than parental MDA-MB-436 cells at each concentration of saruparib (Figure 5I), suggesting that resistance does not arise from the loss of PARP1 trapping. Thus, saruparib treatment of 436-SR-4 and 436-SR-5 cells increases MAR/PARylation and decreases PARP1 trapping. Interestingly, these results are similar to a recent report of a PARP1 L777P mutation that exhibited a similar PARP1 trapping phenotype characterized by release upon titration with saruparib [19].

436-SR-9 and 436-SR-10 cells responded to saruparib treatment nearly identically to one another and in a similar manner to 436-SR-1 cells; both cell lines exhibit only modest saruparib-dependent inhibition of MAR/PARylation and no induction of PARP1 trapping (Figure 5F-I).

Taking these results collectively, the 436-SR-1, 436-SR-9, and 436-SR-10 cell lines exhibit a loss of both saruparib-dependent MAR/PARylation inhibition and PARP1 trapping, either of which may be individually or collectively responsible for saruparib resistance. Intriguingly, 436-SR-4 and 436-SR-5 cells display an overall elevated level of PARP1 trapping, but it is reduced upon saruparib treatment, and MAR/PARylation inhibition is enhanced by saruparib. In addition, these results do not clearly delineate PARP trapping or PARylation inhibition as the driver of PARPi toxicity.

As the SR cell lines exhibited differential sensitivity to the potent PARP trapper talazoparib (Figure 2E), we next evaluated whether these changes in sensitivity may potentially derive from alterations in MAR/PARylation or PARP trapping (Figure S4E-J). 436-SR-1 cells are more sensitive to talazoparib inhibition of MAR/PARylation and exhibit increased PARP1 trapping (Figure S4E-F, K-L), both likely contributing to the increased cellular sensitivity to talazoparib in 436-SR-1 cells (Figure 2E). 436-SR-4 and 436-SR-5 cells, which are also more sensitive to talazoparib (Figure 2E), exhibit resistance to inhibition of MAR/PARylation, however display increased PARP1 trapping by talazoparib as compared to parental MDA-MB-436 cells (Figure S4G-H, K-L). Conversely, 436-SR-9 and 436-SR-10 cells, which are more resistant to talazoparib, exhibit increased inhibition of MAR/PARylation, and comparable and reduced PARP1 trapping, respectively, compared to MDA-MB-436 cells (Figure S4I-L). Taking the previous two results together, the observed changes in inhibition of MAR/PARylation by talazoparib are opposite to the changes in cell survival relative to MDA-MB-436 cells. Instead, PARP1 trapping correlates to cellular sensitivity to talazoparib, arguing that PARP1 trapping by talazoparib is the more toxic event in this case.

### Alternative therapeutic strategies to overcome saruparib resistance

As the emergence of PARPi resistance is a major clinical burden, we evaluated alternative therapeutic strategies that may overcome acquired saruparib resistance. Other characterized mechanisms of PARPi resistance, such as the restoration of HR, result in additional resistance to cisplatin [23]. We found that 436-SR-1 cells exhibited comparable cisplatin sensitivity, while the remaining four cell lines exhibited slight resistance, though significantly less resistance than is observed in cells with the aforementioned PARPi resistance mechanisms (Figure 6A). We next examined SR cell line sensitivity to common chemotherapies for breast cancer patients at various disease stages: the topoisomerase 1 (TOP1) inhibitor camptothecin (CPT) and the tubulin-binding cell division inhibitor paclitaxel. All SR cell lines show decreased sensitivity with 436-SR-1 ∼2-fold, 436-SR-4 and 436-SR-5 ∼7-fold, and 436-SR-9 and 436-SR-10 ∼17-fold increases in CPT resistance (Figure 6B). SR cell lines did not exhibit a significant change in sensitivity to paclitaxel as measured by IC50, but all but 436-SR-1 cells have increased cell viability at the highest paclitaxel concentrations (Figure 6C), likely a reflection of slower growth kinetics and thus fewer cell divisions to be impacted by the cell division inhibitor (Figure 1C). Lastly, we assessed the effectiveness of DDR-targeted agents related to replication stress that have demonstrated anti-cancer activity in contexts of other PARPi resistance mechanisms: the ATR inhibitor ceralasertib, the Wee1 inhibitor MK-1775, and the RPA inhibitor NERx-329 [33–35,41–43]. SR cell lines retained parental-like sensitivity to all three therapeutics, with the general trend that SR cell lines were slightly more sensitive to each of the three DDR inhibitors (Figure 6D-F). This could be indicative of alterations in basal PARP1 activity in the SR cell lines that lead to increased replication stress, bringing about small changes in DDR inhibitor sensitivity.

**Figure 6.**
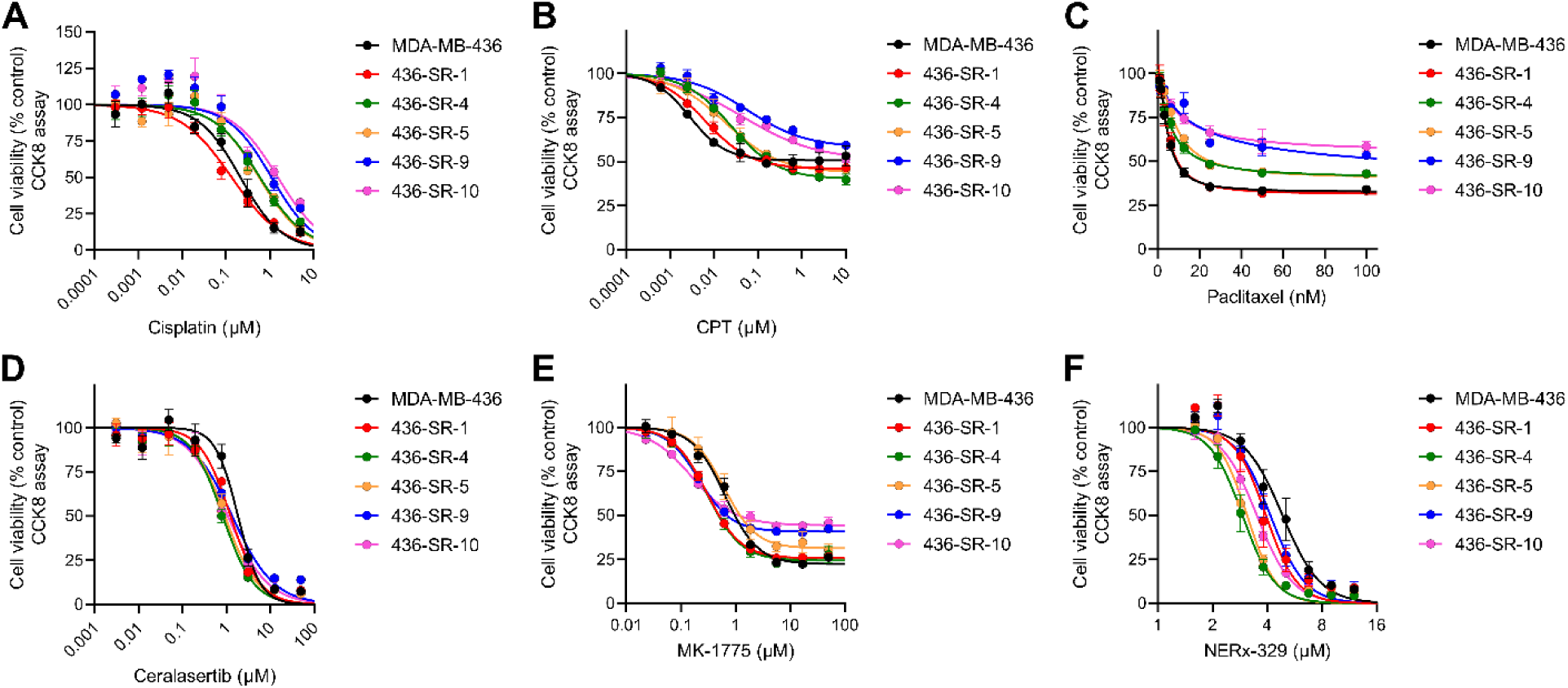
Alternative therapeutic strategies to overcome saruparib resistance. Dose-response curves for MDA-MB-436 and SR cells treated with **A)** cisplatin, **B)** camptothecin (CPT), **C)** paclitaxel, **D)** ceralasertib, **E)** MK-1775, and **F)** NERx-329. All data are depicted as mean ± SEM from triplicate experiments.

## Discussion

PARPi therapy has provided significant increases in survival for some BRCA/HR-deficient cancer patients, while intrinsic and acquired PARPi resistance remains a major clinical limitation. Delineating resistance mechanisms is crucial for designing effective strategies to prevent or overcome resistance. The next generation PARPi saruparib specifically targets PARP1 and is expected to be the front-line PARPi upon completion of ongoing phase 3 clinical trials, and though it has demonstrated effective and durable responses, saruparib resistance will undoubtedly be a barrier to widespread utility towards curative intent. The degree to which currently established pan-PARPi resistance mechanisms are relevant in cases of acquired saruparib resistance is currently unknown, and there may well be novel and distinct saruparib resistance mechanisms due to the differences in selectivity. To address these possibilities, we generated and characterized five saruparib resistant cell lines and identified three PARP1 catalytic domain mutations that drive saruparib resistance. Strikingly, SR cell lines do not exhibit pan-PARPi resistance, with each PARP1 mutation conferring differential sensitivity to each PARPi examined, differing from other PARPi resistant cell lines generated via treatment with rucaparib or olaparib [23,36]. Therefore, patients who develop PARPi resistance due to G871W or K893E PARP1 mutations, for example, may well benefit from treatment from an additional PARPi, such as olaparib or talazoparib, while those with the D770H PARP1 mutation would benefit from alternative therapeutic strategies. Importantly, platinum-based agents remain viable treatment options for saruparib resistance, differing from other pan-PARPi resistance mechanisms like HR restoration that are resistant to platinum-based therapies [23]. Moreover, our work supports the use of common breast cancer therapeutics like taxanes and TOP1 inhibitors in treating saruparib-resistant cancers, coinciding with the recent surge of TOP1 antibody-drug conjugate strategies [44].

Of our five SR cell lines, all possess PARP1 catalytic domain mutations that are driving resistance. Although these mutations confer differences in PARP1 catalytic activity and trapping, this represents a unifying mechanism of saruparib resistance. During the preparation of this manuscript, an independent report that is to our knowledge the only other known characterization of a specific mechanism of acquired saruparib resistance identified another PARP1 catalytic domain mutation (L777P) that confers resistance to saruparib in a different BRCA1-deficient breast cancer cell line (SUM149PT) [19]. This mutation appears to function like the K893E mutation in 436-SR-4 and 436-SR-5 cells in terms of PARP1 trapping, and L777P PARP1-containing cells were similarly more sensitive to olaparib [19]. However, these cells were not more sensitive to talazoparib while 436-SR-4 and 436-SR-5 cells exhibited a ∼10-fold increase in talazoparib efficacy over parental MDA-MB-436 cells. Taking this result into consideration with the two additional mutations from this study (G871W and D770H), there are currently seven characterized cell lines with acquired saruparib resistance, and in all cases, resistance is driven by one of four PARP1 catalytic domain mutations [19]. Therefore, PARP1 mutation are likely to be a major driver of saruparib resistance due to its PARP1-selective nature. Although we await genomic data from patients who progress on saruparib to determine if this mechanism uncovered in cell culture also holds in the clinic, this work suggests the exciting possibility that acquired saruparib resistance is treatable via other clinical PARPi depending upon the specific PARP1 mutation. The identification of PARP1 mutations that confer saruparib resistance and characterization of their sensitivity to other clinical PARPi will be key in the treatment of saruparib resistant cancers. Moreover, the retained sensitivity to other common breast cancer or DDR-targeted therapeutics are additional therapeutic alternatives. Therefore, along with the potency of saruparib in treating BRCA/HR-deficient cancers and the significantly improved safety profile, the PARP1 selectivity of saruparib may select for PARP1 mutations as the primary mechanism of resistance that ultimately renders the treatment of saruparib resistant cancers much more manageable than current clinical pan-PARPi resistant cancers.

## Acknowledgements

We thank the IU Melvin and Bren Simon Comprehensive Cancer Center, IU Medical Genomics Core, IU Cellular Response Technologies Core, and Dr. Emma Doud and the IU Center for Proteome Analysis for their support. We would also like to thank Dr. Lata Balakrishnan for providing the PARP-CAT expression plasmid and Eeson Rajendra for providing the long-template HR plasmid.

## Data Availability

All primary data from this study are included in the main text and supporting information.

## Author Contributions

M.R.J. and J.J.T. conceived and designed the study. M.R.J., J.L.K., and J.E.G. performed experiments. S.L. and J.W. performed bioinformatic analyses. All authors analyzed and interpreted the results and contributed to the writing and preparation of the manuscript.

## Materials and Methods

### Cell Culture

MDA-MB-436 cells (ATCC) were cultured in 1:1 DMEM:Ham’s F12 medium (Corning) supplemented with 10% FBS and penicillin/streptomycin. After saruparib-resistant cell lines were established, they were maintained in the same medium supplemented with 3 nM saruparib. All cells were grown at 37°C with 5% CO_2_.

### Generation of Saruparib-Resistant Cell Lines

2.2×10^7^ MDA-MB-436 cells were plated at 2×10^6^ cells/plate in 150 mm dishes. Cells were allowed to grow an additional 2 days and double to an estimated ∼5×10^7^ cells total before applying saruparib treatments. Three treatment regimes were used (see Figure S1A) applying alternating 1 week treatment/1 week recovery, 2 weeks treatment/1 week recovery, or 3 weeks treatment/1 week recovery with 30 nM saruparib. Recovery medium/treatments were refreshed twice per week and PBS wash was performed twice per week to remove cell debris. Treatments were continued for ∼5 weeks before colonies emerged. Colonies were isolated by using cloning disks soaked in trypsin and each colony was transferred to a well of a 24-well plate. Isolated clones that exhibited sustained growth to confluence in the 24-well plate were transferred to larger flasks for expansion and cell stocks were frozen in liquid nitrogen. Isolated saruparib-resistant cell lines were grown in medium supplemented with 3 nM saruparib as a maintenance pressure.

### CCK-8 Cell Viability Assays

For camptothecin, MK-1775, NERx-329, and paclitaxel treatments (inhibitor stocks dissolved in DMSO), cells were plated at 5×10^3^ cells/well in a 96-well plate and incubated for 16-24 hours before treatments. Treatments were performed at the indicated concentrations or vehicle control (0.5% DMSO final concentration) for 48 hours before cell viability was assessed using the CCK-8 assay (see below). For cisplatin treatments (cisplatin stock dissolved in sterile water), cells were plated at 2.5×10^3^ cells/well in a 96-well plate and incubated for 16-24 hours before treatments. Treatments were performed at the indicated concentrations or vehicle control for 120 hours before cell viability was assessed using the CCK-8 assay (see below). For ceralasertib, olaparib, niraparib, saruparib, stenoparib, and talazoparib treatments (inhibitor stocks dissolved in DMSO), cells were plated at 2.5×10^3^ cells/well in a 96-well plate and incubated for 16-24 hours before treatments. Treatments were performed at the indicated concentrations or vehicle control (0.5% DMSO final concentration) for 144 hours before cell viability was assessed using the CCK-8 assay. 10 µL of CCK-8 reagent was added to each well and incubated 1-2 hours. Absorbance at 450 nm was measured using a BioTek Synergy H1 plate reader and values were normalized to vehicle control wells. All cell viability assays were performed in triplicate.

### In vivo Analysis

Female NOD-scid/IL2Rg-null (NSG) mice (*In Vivo* Therapeutics Core Facility, IU Simon Comprehensive Cancer Center) were housed in a pathogen-free facility at the IUSM LARC facility. Animal studies were approved by the Institutional Animal Care and Use Committee of the Indiana University School of Medicine. The hind flanks of 8-10-week-old NSG mice were implanted with MDA-MB-436 or 436-SR-1 cells (∼4×10^6^) in Matrigel. Tumors volume was monitored using electronic calipers [tumor volume = length x (perpendicular width)2 x 0.5]. Tumors were allowed to grow to an average of ∼325 mm^3^ and mice were randomized into vehicle or treatment arms (n=7 and n=6 for vehicle and treatment arms, respectively, for both cell lines). Saruparib was dissolved in acidic water (pH = 2-3) and administered via oral gavage at 0.3 mg/kg daily following a 5 days on-2 days off treatment cycle. Tumor volumes were monitored over time and the final tumor weight was determined from the excised tumors at the end of the experimental period.

### Live-Cell Imaging

Cells were plated at 2.5×10^3^ cells/well in a 96-well plate and incubated for 7 days in an Incucyte imager. Images were captured at 4x every 4 hours and confluence was determined using Incucyte software applying a specific mask designed for MDA-MB-436 cells. Confluence was normalized according to the initial percent confluence and a final percent confluence of 100%.

### Homologous Recombination Reporter Assay

Homologous recombination DNA repair activity was measured via an extrachromosomal reporter assay according to previously published literature with slight modifications [45]. Briefly, cell lines were reverse transfected with the long-template HR repair specific reporter (1 µg DNA/1 x 10^6^ cells) and transfection control plasmid (0.66 µg DNA /1 x 10^6^ cells) using JetPRIME® in a 1:3 DNA to transfection reagent ratio. Transfected cells were plated 30,000 cell/well and incubated for 24 hours. Post incubation, LT HR repair was measured and quantified using the Nano-Glo® Dual-Luciferase® Reporter Assay System according to the manufacturer’s protocol.

### Whole Genome Sequencing

2×10^6^ cells from each cell line were collected and genomic DNA was isolated using the DNEasy Blood and Tissue Kit (Qiagen). DNA quality was confirmed using the Bioanalyzer system (Agilent). FASTQ files were aligned to the human reference genome (hg38) using BWA (v. 0.7.17, http://bio-bwa.sourceforge.net/) [46]. PCR duplicates were removed and coverage metrics were calculated using PICARD-tools through The Genome Analysis Toolkit (GATK, v. 4.2.6.1, http://www.broadinstitute.org/gatk/) [47]. GATK was used for SNP and INDEL discovery according to GATK best practices [48]. ANNOVAR (https://annovar.openbioinformatics.org/) was used to annotate resulting variants [49]. Panel of normal somatic-hg38_1000g_pon.hg38.vcf.gz and MDA-MB-436_S2 were used as normal sample.

### RNA-Seq

0.1-1.0×10^7^ cells from each cell line were collected and RNA was isolated using the RNEasy Mini Kit (Qiagen). DNA quality was confirmed using the Bioanalyzer system (Agilent). The reads were mapped to the human genome hg38 using STAR (v2.7.11a) [50]. Uniquely mapped sequencing reads were assigned to GENCODE G47 gene using featureCounts [51] with the following parameters: “-s 2 –p –Q 10 - O”. The data was filtered using read count > 10 in at least one of the samples, normalized using TMM (trimmed mean of M values) method and subjected to differential expression analysis using edgeR [52,53]. Gene ontology and KEGG pathway functional analysis was performed on differential expression gene with false discovery rate cut-off of 0.05 and absolute value of log2 of fold change cut-off of 1 using DAVID [54,55]. Allele-Specific Expression of PARP1 was derived using GATK ASEReadCounter.

### Purification of PARP1-CAT and Mutants

The PARP1-CAT plasmid (PARP1 661-1014 L713F) containing a thrombin-cleavable C-terminal His_6_-tag was kindly provided by Dr. Lata Balakrishnan. The thrombin cleavage site was converted into a TEV cleavage site using standard cloning procedures and verified via DNA sequencing. The D770H, G871W, and K893E mutations were inserted using standard mutagenesis procedures and verified via DNA sequencing. PARP1-CAT and the three mutants were all purified in the same manner as follows. Plasmids were individually transformed into *E. coli* BL21 cells and plated on LB plates supplemented with 50 µg/mL kanamycin. Precultures derived from a single colony were inoculated into 0.5 L LB supplemented with 50 µg/mL kanamycin and 10 mM benzamide (to inhibit PARP1 activity) and grown at 37°C with shaking. Once OD_600_ = 0.6-0.8, protein expression was induced with the addition of 0.2 mM IPTG and cells were grown for an additional ∼16 hours at 18°C. Cells were collected by centrifugation and resuspended in 25 mM Tris pH 8, 500 mM NaCl, 10 mM imidazole, 2 mM BME, 0.1% NP-40, 0.5 mg/mL lysozyme, 0.5 mM PMSF, 1 µg/mL pepstatin A, and 1 µg/mL leupeptin. Cells were lysed by Dounce homogenizer and sonication for 15 min. Cell extract was separated by centrifugation at 11,000 x g for 20 min at 4°C and loaded onto a Ni-NTA column pre-equilibrated with Ni Buf A (25 mM Tris pH 8, 500 mM NaCl, 10 mM imidazole, 2 mM BME). The column was then washed with 5 column volumes (CV) Ni Buf A and the PARP-CAT-His_6_ was eluted with 5 CV Ni Buf B (25 mM Tris pH 8, 500 mM NaCl, 400 mM imidazole, 2 mM BME). To the elution, 1 mM DTT and 1,000 IU His-TEV (Neta Scientific) was added and incubated for 2 hours at room temperature with occasional stirring. The elution was then dialyzed for 16-18 hr at 4°C into 25 mM Tris pH 8, 500 mM NaCl, 2 mM BME. The dialyzed sample was re-loaded onto a Ni-NTA column pre-equilibrated with Ni Buf A. The flow-through containing the cleaved PARP1-CAT was collected and fractions containing PARP1-CAT at >95% purity as assessed by SDS-PAGE were combined, aliquoted, snap frozen in liquid nitrogen, and stored at −80°C. Each mutation was confirmed by the IU School of Medicine Proteomics Core.

### PARP1-CAT Auto-PARylation Assay

Reactions were conducted in reaction buffer (20 mM Tris pH 7.5, 50 mM NaCl, 5 mM MgCl_2_, 0.1 mM DTT, and 0.5% DMSO). 50 nM PARP1-CAT or mutant were incubated in reaction buffer with or without saruparib/talazoparib at 25°C for 10 min. Reactions were initiated with the addition of 250 µM NAD^+^ and conducted for 5 and 15 min (time-course experiments) or 15 min (PARPi experiments) at 25°C. Reactions were quenched by the addition of 6x quenching dye (250 mM Tris pH 6.8, 24% glycerol, 8% SDS, 0.12% bromophenol blue, 100 mM EDTA) and briefly incubate at 95°C. To detect auto-PARylation via Western blot, samples were run on SDS-PAGE and transferred to a polyvinylidene difluoride (PVDF) membranes. Membranes were blocked with 2% BSA in Tris buffered-saline, 0.5% Tween 20 (TBST) for 2 hr at room temperature before incubation with MAR/PAR (E6F6A) primary antibody (Cell Signaling Technology, 1:10,000 in 2% BSA in TBST) at 4°C overnight. Membranes were washed with TBST and incubated with goat anti-rabbit IgG-horseradish peroxidase (HRP) conjugate secondary antibody (Bio-Rad) in 2% BSA in TBST for 2 hr at room temperature. Membranes were washed with TBST and immunoblots were detected using the chemiluminescent Clarity Western ECL Substrate (Bio-Rad).

### Soluble and Chromatin-Bound Extracts for PARP Trapping Analysis

Cells were plated at 1×10^6^ cells/well in a 6-well plate and incubated for 16-24 hours before treatments. Cells were treated with vehicle DMSO (kept at 0.5% for all treatments), 0.1 mg/mL methyl methanesulfonate (prepared fresh), and/or 0.2 nM, 2 nM, 20 nM or 200 nM saruparib or talazoparib at 37°C. After 2 hr, cells were quickly washed with PBS and trypsinized before being resuspended in 250 µL lysis buffer (50 mM HEPES pH 7.4, 100 mM KCl, 2.5 mM MgCl_2_, 5 mM EDTA, 3 mM DTT, 0.5% Triton X-100, 10% glycerol, and 1x Halt Protease and Phosphatase Inhibitor Cocktail [ThermoFisher Scientific]). The mixture was incubated on ice for 45 minutes and soluble and chromatin-bound fractions were by centrifugation at 16,000 xg for 15 minutes at 4°C. The supernatant was collected as the soluble fraction and the pellet containing the chromatin-bound fraction was washed twice more with 250 µL lysis buffer. The pellet was then resuspended in 250 µL lysis buffer. The protein concentration was determined for the soluble fractions by Pierce BCA Protein assay (Fisher Scientific) and equivolumes of soluble and chromatin-bound fractions from each sample were used to load the equivalent of 15 µg of soluble protein for SDS-PAGE separation. Proteins were then transferred to PVDF membranes and blocked in 2% BSA in TBST for 1 hr at room temperature before incubating with primary antibody overnight at 4°C. The primary antibodies used were PARP1 (Cell Signaling Technology) at 1:10,000, PARP2 (F3Z6Z, Cell Signaling Technology) at 1:5,000, MAR/PAR (E6F6A, Cell Signaling Technology) at 1:10,000, GAPDH (GA1R, Invitrogen) at 1:10:000, and Histone H3 (Abcam) at 1:10,000. Membranes were then washed with TBST and incubated with goat anti-rabbit IgG-HRP or goat anti-mouse IgG-HRP secondary antibody (Bio-Rad) 1:5,000 in 2% BSA in TBST for 2 hr at room temperature. Membranes were washed with TBST and immunoblots were detected using the chemiluminescent Clarity Western ECL Substrate (Bio-Rad).

## References

1. Petrucelli N, Daly MB, Pal T. BRCA1- and BRCA2-Associated Hereditary Breast and Ovarian Cancer. Seattle (WA): University of Washington, Seattle; 1993.

2. Farmer H, McCabe N, Lord CJ, Tutt ANJ, Johnson DA, Richardson TB, et al. Targeting the DNA repair defect in BRCA mutant cells as a therapeutic strategy. Nature. 2005;434:917–21. 10.1038/nature03445

3. Bryant HE, Schultz N, Thomas HD, Parker KM, Flower D, Lopez E, et al. Specific killing of BRCA2-deficient tumours with inhibitors of poly(ADP-ribose) polymerase. Nature. 2005;434:913–7. 10.1038/nature03443

4. Wicks AJ, Krastev DB, Pettitt SJ, Tutt ANJ, Lord CJ. Opinion: PARP inhibitors in cancer-what do we still need to know? Open Biol. 2022;12:220118. 10.1098/rsob.220118

5. Suskiewicz MJ, Munnur D, Strømland Ø, Yang J-C, Easton LE, Chatrin C, et al. Updated protein domain annotation of the PARP protein family sheds new light on biological function. Nucleic Acids Res. 2023;51:8217–36. 10.1093/nar/gkad514

6. Shieh WM, Amé J-C, Wilson M V., Wang Z-Q, Koh DW, Jacobson MK, et al. Poly(ADP-ribose) Polymerase Null Mouse Cells Synthesize ADP-ribose Polymers. Journal of Biological Chemistry. 1998;273:30069–72. 10.1074/jbc.273.46.30069

7. Amé J-C, Rolli V, Schreiber V, Niedergang C, Apiou F, Decker P, et al. PARP-2, A Novel Mammalian DNA Damage-dependent Poly(ADP-ribose) Polymerase. Journal of Biological Chemistry. 1999;274:17860–8. 10.1074/jbc.274.25.17860

8. Langelier M-F, Planck JL, Roy S, Pascal JM. Structural Basis for DNA Damage–Dependent Poly(ADP-ribosyl)ation by Human PARP-1. Science (1979). 2012;336:728–32. 10.1126/science.1216338

9. Dawicki-McKenna JM, Langelier M-F, DeNizio JE, Riccio AA, Cao CD, Karch KR, et al. PARP-1 Activation Requires Local Unfolding of an Autoinhibitory Domain. Mol Cell. 2015;60:755–68. 10.1016/j.molcel.2015.10.013

10. Gu Kang B, Kang S-U, Jin Kim J, Kwon J-S, Gagné J-P, Yun Lee S, et al. Proteome-wide microarray-based screening of PAR-binding proteins. Nucleic Acids Res. 2025;53. 10.1093/nar/gkaf300

11. Helleday T. The underlying mechanism for the PARP and BRCA synthetic lethality: clearing up the misunderstandings. Mol Oncol. 2011;5:387–93. 10.1016/j.molonc.2011.07.001

12. Patel AG, Sarkaria JN, Kaufmann SH. Nonhomologous end joining drives poly(ADP-ribose) polymerase (PARP) inhibitor lethality in homologous recombination-deficient cells. Proceedings of the National Academy of Sciences. 2011;108:3406–11. 10.1073/pnas.1013715108

13. Schlacher K, Christ N, Siaud N, Egashira A, Wu H, Jasin M. Double-Strand Break Repair-Independent Role for BRCA2 in Blocking Stalled Replication Fork Degradation by MRE11. Cell [Internet]. Elsevier B.V.; 2011;145:529–42. 10.1016/j.cell.2011.03.041

14. Ray Chaudhuri A, Callen E, Ding X, Gogola E, Duarte AA, Lee J-E, et al. Replication fork stability confers chemoresistance in BRCA-deficient cells. Nature. Nature Publishing Group; 2016;535:382–7. 10.1038/nature18325

15. Panzarino NJ, Krais JJ, Cong K, Peng M, Mosqueda M, Nayak SU, et al. Replication Gaps Underlie BRCA Deficiency and Therapy Response. Cancer Res. American Association for Cancer Research Inc.; 2021;81:1388–97. 10.1158/0008-5472.CAN-20-1602

16. Cong K, Peng M, Kousholt AN, Lee WTC, Lee S, Nayak S, et al. Replication gaps are a key determinant of PARP inhibitor synthetic lethality with BRCA deficiency. Mol Cell. Cell Press; 2021;81:3128–3144.e7. 10.1016/j.molcel.2021.06.011

17. Murai J, Huang SN, Das BB, Renaud A, Zhang Y, Doroshow JH, et al. Trapping of PARP1 and PARP2 by Clinical PARP Inhibitors. Cancer Res. 2012;72:5588–99. 10.1158/0008-5472.CAN-12-2753

18. Pommier Y, O’Connor MJ, de Bono J. Laying a trap to kill cancer cells: PARP inhibitors and their mechanisms of action. Sci Transl Med. 2016;8. 10.1126/scitranslmed.aaf9246

19. Ribeiro J, Hansen K, van Beek L, Stubbs C, Frigola J, Talbot S, et al. Genetic evidence for PARP1 trapping as a driver of PARP inhibitor efficacy in *BRCA* mutant cancer cells. Nucleic Acids Res. 2025;53. 10.1093/nar/gkaf1398

20. MacGilvary N, Cantor SB. Positioning loss of PARP1 activity as the central toxic event in BRCA-deficient cancer. DNA Repair (Amst). 2024;144:103775. 10.1016/j.dnarep.2024.103775

21. Petropoulos M, Karamichali A, Rossetti GG, Freudenmann A, Iacovino LG, Dionellis VS, et al. Transcription–replication conflicts underlie sensitivity to PARP inhibitors. Nature. 2024;628:433–41. 10.1038/s41586-024-07217-2

22. Zou Y, Zhang H, Chen P, Tang J, Yang S, Nicot C, et al. Clinical approaches to overcome PARP inhibitor resistance. Mol Cancer. 2025;24:156. 10.1186/s12943-025-02355-1

23. Johnson N, Johnson SF, Yao W, Li Y-C, Choi Y-E, Bernhardy AJ, et al. Stabilization of mutant BRCA1 protein confers PARP inhibitor and platinum resistance. Proc Natl Acad Sci U S A. 2013;110:17041–6. 10.1073/pnas.1305170110

24. Rogers CM, Kaur H, Swift ML, Raina VB, Zhou S, Kawale AS, et al. CTC1-STN1-TEN1 controls DNA break repair pathway choice via DNA end resection blockade. Science. 2025;388:881–8. 10.1126/science.adt3034

25. Nesic K, Kondrashova O, Hurley RM, McGehee CD, Vandenberg CJ, Ho G-Y, et al. Acquired RAD51C Promoter Methylation Loss Causes PARP Inhibitor Resistance in High-Grade Serous Ovarian Carcinoma. Cancer Res. 2021;81:4709–22. 10.1158/0008-5472.CAN-21-0774

26. Teng Q-X, Lei Z-N, Wang J-Q, Yang Y, Wu Z-X, Acharekar ND, et al. Overexpression of ABCC1 and ABCG2 confers resistance to talazoparib, a poly (ADP-Ribose) polymerase inhibitor. Drug Resistance Updates. 2024;73:101028. 10.1016/j.drup.2023.101028

27. Pettitt SJ, Krastev DB, Brandsma I, Dréan A, Song F, Aleksandrov R, et al. Genome-wide and high-density CRISPR-Cas9 screens identify point mutations in PARP1 causing PARP inhibitor resistance. Nat Commun. 2018;9:1849. 10.1038/s41467-018-03917-2

28. Gogola E, Duarte AA, de Ruiter JR, Wiegant WW, Schmid JA, de Bruijn R, et al. Selective Loss of PARG Restores PARylation and Counteracts PARP Inhibitor-Mediated Synthetic Lethality. Cancer Cell. 2018;33:1078–1093.e12. 10.1016/j.ccell.2018.05.008

29. Farrés J, Martín-Caballero J, Martínez C, Lozano JJ, Llacuna L, Ampurdanés C, et al. Parp-2 is required to maintain hematopoiesis following sublethal γ-irradiation in mice. Blood. 2013;122:44–54. 10.1182/blood-2012-12-472845

30. Farrés J, Llacuna L, Martin-Caballero J, Martínez C, Lozano JJ, Ampurdanés C, et al. PARP-2 sustains erythropoiesis in mice by limiting replicative stress in erythroid progenitors. Cell Death Differ. 2015;22:1144–57. 10.1038/cdd.2014.202

31. AZD5305 More Tolerable than Earlier PARP Agents. Cancer Discov. 2022;12:1602–1602. 10.1158/2159-8290.CD-NB2022-0039

32. Illuzzi G, Staniszewska AD, Gill SJ, Pike A, McWilliams L, Critchlow SE, et al. Preclinical Characterization of AZD5305, A Next-Generation, Highly Selective PARP1 Inhibitor and Trapper. Clinical Cancer Research. 2022;28:4724–36. 10.1158/1078-0432.CCR-22-0301

33. Herencia-Ropero A, Llop-Guevara A, Staniszewska AD, Domènech-Vivó J, García-Galea E, Moles-Fernández A, et al. The PARP1 selective inhibitor saruparib (AZD5305) elicits potent and durable antitumor activity in patient-derived BRCA1/2-associated cancer models. Genome Med. 2024;16:107. 10.1186/s13073-024-01370-z

34. Arce-Gallego S, Esquefa V, Domenech H, Salca A, Aguilar D, Casals T, et al. Combination therapy overcomes secondary PARPi resistance in ATM-deficient prostate cancer. NPJ Precis Oncol. 2025;9:407. 10.1038/s41698-025-01179-y

35. Yazinski SA, Comaills V, Buisson R, Genois M-M, Nguyen HD, Ho CK, et al. ATR inhibition disrupts rewired homologous recombination and fork protection pathways in PARP inhibitor-resistant BRCA-deficient cancer cells. Genes Dev. 2017;31:318–32. 10.1101/gad.290957.116

36. Shih C-T, Huang T-T, Nair JR, Ibanez KR, Lee J-M. Poly (ADP-Ribose) Polymerase Inhibitor Olaparib-Resistant BRCA1-Mutant Ovarian Cancer Cells Demonstrate Differential Sensitivity to PARP Inhibitor Rechallenge. Cells. 2024;13:1847. 10.3390/cells13221847

37. Ooi S-K, Sato S, Tomomori-Sato C, Zhang Y, Wen Z, Banks CAS, et al. Multiple roles for PARP1 in ALC1-dependent nucleosome remodeling. Proceedings of the National Academy of Sciences. 2021;118. 10.1073/pnas.2107277118

38. Cortes U, Tong W-M, Coyle DL, Meyer-Ficca ML, Meyer RG, Petrilli V, et al. Depletion of the 110-Kilodalton Isoform of Poly(ADP-Ribose) Glycohydrolase Increases Sensitivity to Genotoxic and Endotoxic Stress in Mice. Mol Cell Biol. 2004;24:7163–78. 10.1128/MCB.24.16.7163-7178.2004

39. Haince J-F, McDonald D, Rodrigue A, Déry U, Masson J-Y, Hendzel MJ, et al. PARP1-dependent kinetics of recruitment of MRE11 and NBS1 proteins to multiple DNA damage sites. J Biol Chem. 2008;283:1197–208. 10.1074/jbc.M706734200

40. Kanev P-B, Varhoshkova S, Georgieva I, Lukarska M, Kirova D, Danovski G, et al. A unified mechanism for PARP inhibitor-induced PARP1 chromatin retention at DNA damage sites in living cells. Cell Rep. 2024;43:114234. 10.1016/j.celrep.2024.114234

41. Chiappa M, Guffanti F, Anselmi M, Lupi M, Panini N, Wiesmüller L, et al. Combinations of ATR, Chk1 and Wee1 Inhibitors with Olaparib Are Active in Olaparib Resistant Brca1 Proficient and Deficient Murine Ovarian Cells. Cancers (Basel). 2022;14:1807. 10.3390/cancers14071807

42. VanderVere-Carozza PS, Jordan MR, Garrett JE, Pollok KE, Pawelczak KS, Turchi JJ. Replication protein A protects lagging strand gaps, restricting PARP inhibitor-induced synthetic lethality in BRCA1-deficient tumors. Nucleic Acids Res. 2026;54. 10.1093/nar/gkag396

43. Gupta N, Huang T-T, Horibata S, Lee J-M. Cell cycle checkpoints and beyond: Exploiting the ATR/CHK1/WEE1 pathway for the treatment of PARP inhibitor–resistant cancer. Pharmacol Res. 2022;178:106162. 10.1016/j.phrs.2022.106162

44. Pommier Y, Thomas A. New Life of Topoisomerase I Inhibitors as Antibody–Drug Conjugate Warheads. Clinical Cancer Research. 2023;29:991–3. 10.1158/1078-0432.CCR-22-3640

45. Rajendra E, Grande D, Mason B, Di Marcantonio D, Armstrong L, Hewitt G, et al. Quantitative, titratable and high-throughput reporter assays to measure DNA double strand break repair activity in cells. Nucleic Acids Res. 2024;52:1736–52. 10.1093/nar/gkad1196

46. Li H, Durbin R. Fast and accurate short read alignment with Burrows–Wheeler transform. Bioinformatics. 2009;25:1754–60. 10.1093/bioinformatics/btp324

47. McKenna A, Hanna M, Banks E, Sivachenko A, Cibulskis K, Kernytsky A, et al. The Genome Analysis Toolkit: A MapReduce framework for analyzing next-generation DNA sequencing data. Genome Res. 2010;20:1297–303. 10.1101/gr.107524.110

48. DePristo MA, Banks E, Poplin R, Garimella K V, Maguire JR, Hartl C, et al. A framework for variation discovery and genotyping using next-generation DNA sequencing data. Nat Genet. 2011;43:491–8. 10.1038/ng.806

49. Wang K, Li M, Hakonarson H. ANNOVAR: functional annotation of genetic variants from high-throughput sequencing data. Nucleic Acids Res. 2010;38:e164–e164. 10.1093/nar/gkq603

50. Dobin A, Davis CA, Schlesinger F, Drenkow J, Zaleski C, Jha S, et al. STAR: ultrafast universal RNA-seq aligner. Bioinformatics. 2013;29:15–21. 10.1093/bioinformatics/bts635

51. Liao Y, Smyth GK, Shi W. featureCounts: an efficient general purpose program for assigning sequence reads to genomic features. Bioinformatics. 2014;30:923–30. 10.1093/bioinformatics/btt656

52. Robinson MD, McCarthy DJ, Smyth GK. edgeR : a Bioconductor package for differential expression analysis of digital gene expression data. Bioinformatics. 2010;26:139–40. 10.1093/bioinformatics/btp616

53. McCarthy DJ, Chen Y, Smyth GK. Differential expression analysis of multifactor RNA-Seq experiments with respect to biological variation. Nucleic Acids Res. 2012;40:4288–97. 10.1093/nar/gks042

54. Dennis G, Sherman BT, Hosack DA, Yang J, Gao W, Lane HC, et al. DAVID: Database for Annotation, Visualization, and Integrated Discovery. Genome Biol. 2003;4:P3.

55. Huang DW, Sherman BT, Lempicki RA. Systematic and integrative analysis of large gene lists using DAVID bioinformatics resources. Nat Protoc. 2009;4:44–57. 10.1038/nprot.2008.211

