## Supplementary Information for "PARP1 catalytic domain mutations drive high-level resistance to saruparib while preserving DNA damage response vulnerabilities"

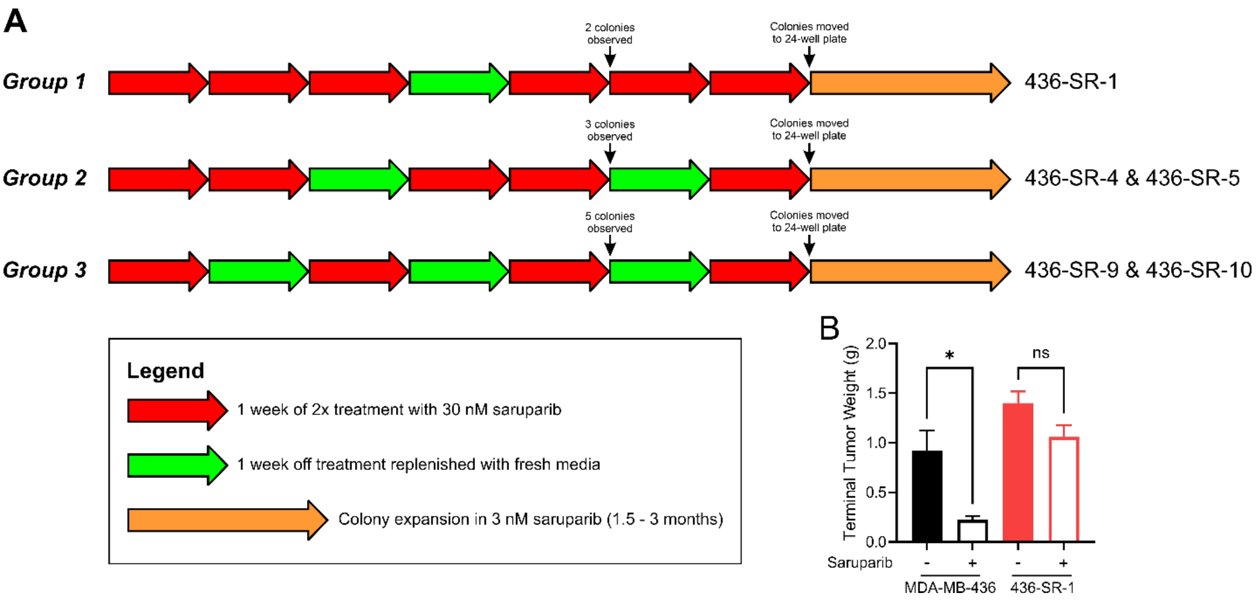


**Figure S1. A)** Depiction of the three treatments used including the timing of colony formation, initiation of colony expansion, and the resulting cell lines that were established. **B)** Terminal tumor weight of MDA-MB-436 and 436-SR-1 tumors treated with vehicle or saruparib from the experiment shown in Figure 1F-G.

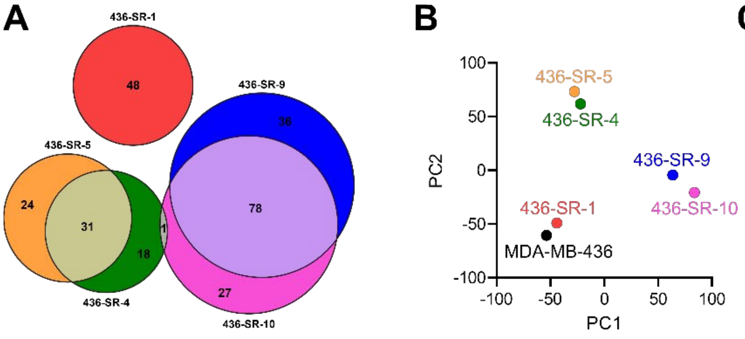


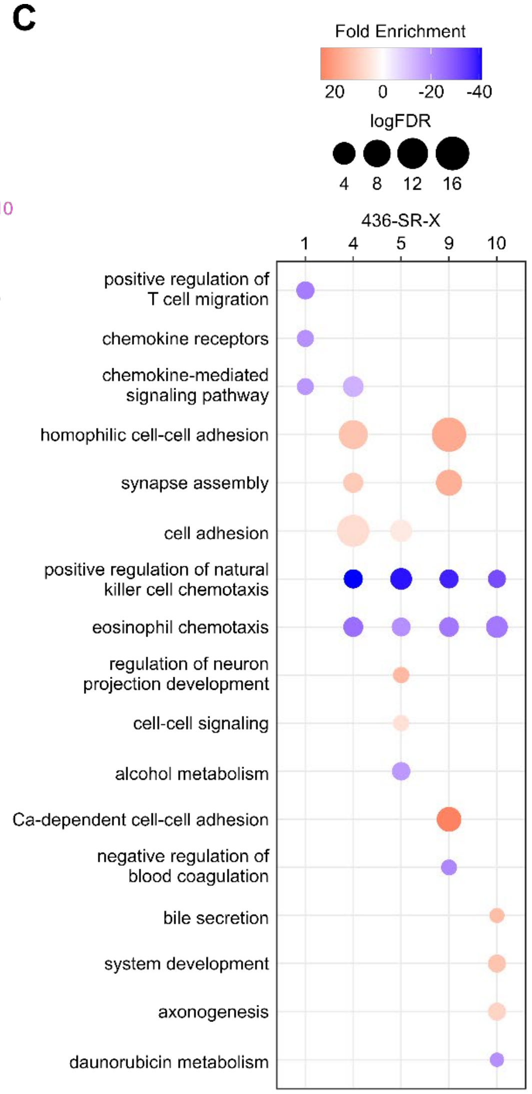


**D**

| | 436-SR-1 (G871W) | 436-SR-4 (K893E) | 436-SR-5 (K893E) | 436-SR-9 (D770H) | 436-SR-10 (D770H) |
|---|---|---|---|---|---|
| **WT Allele Frequency** | 70% | 26% | 52% | 54% | 52% |
| **Mutant Allele Frequency** | 30% | 74% | 48% | 46% | 48% |

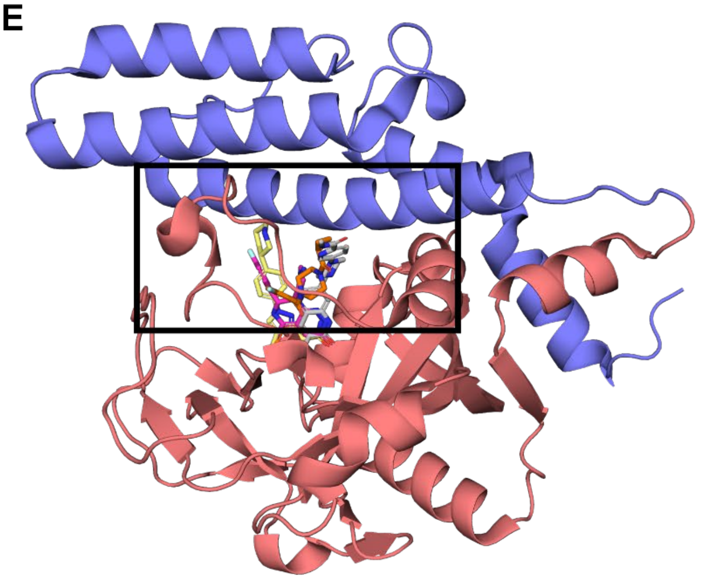


**Figure S2. A)** Number of amino acid altering mutations detected from whole genome sequencing for each SR cell line. **B)** Principal component analysis of RNA-seq data set comparing each SR cell line. **C)** Top three up- and down-regulated pathways from the RNA-seq data set for all SR cell lines relative to the parental MDA-MB-436 cell line. 436-SR-1 cells did not have any significantly upregulated pathways. **D)** Allele-specific expression of wild-type (WT) and mutant alleles for PARP1 transcripts in each SR cell line from the RNA-seq data set. **E)** Structure of the PARP1 catalytic domain bound to saruparib (gray, PDB: 9ETQ), olaparib (orange, PDB: 7AAD), talazoparib (magenta, PDB: 7KK3), and niraparib (yellow, PDB: 7KK5) generated by aligning catalytic domains from each referenced structure to the saruparib-bound structure. The black box depicts the zoomed view of Figure 1C that includes the location of the identified SR cell line PARP1 mutations.

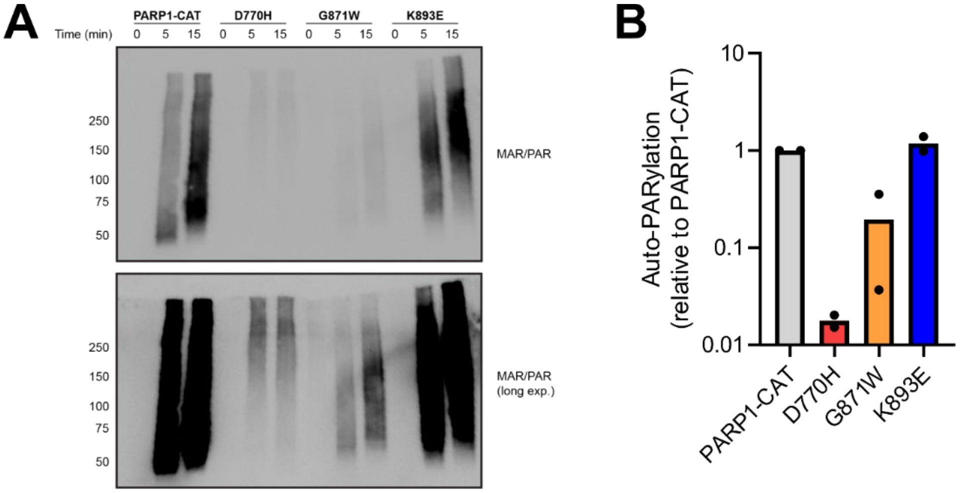


**Figure S3. A)** Representative time-course comparison of auto-PARylation activity of PARP1-CAT and the three SR cell line PARP1 mutations. **B)** Quantification of the time course data from panel A depicted as the mean from duplicate experiments.

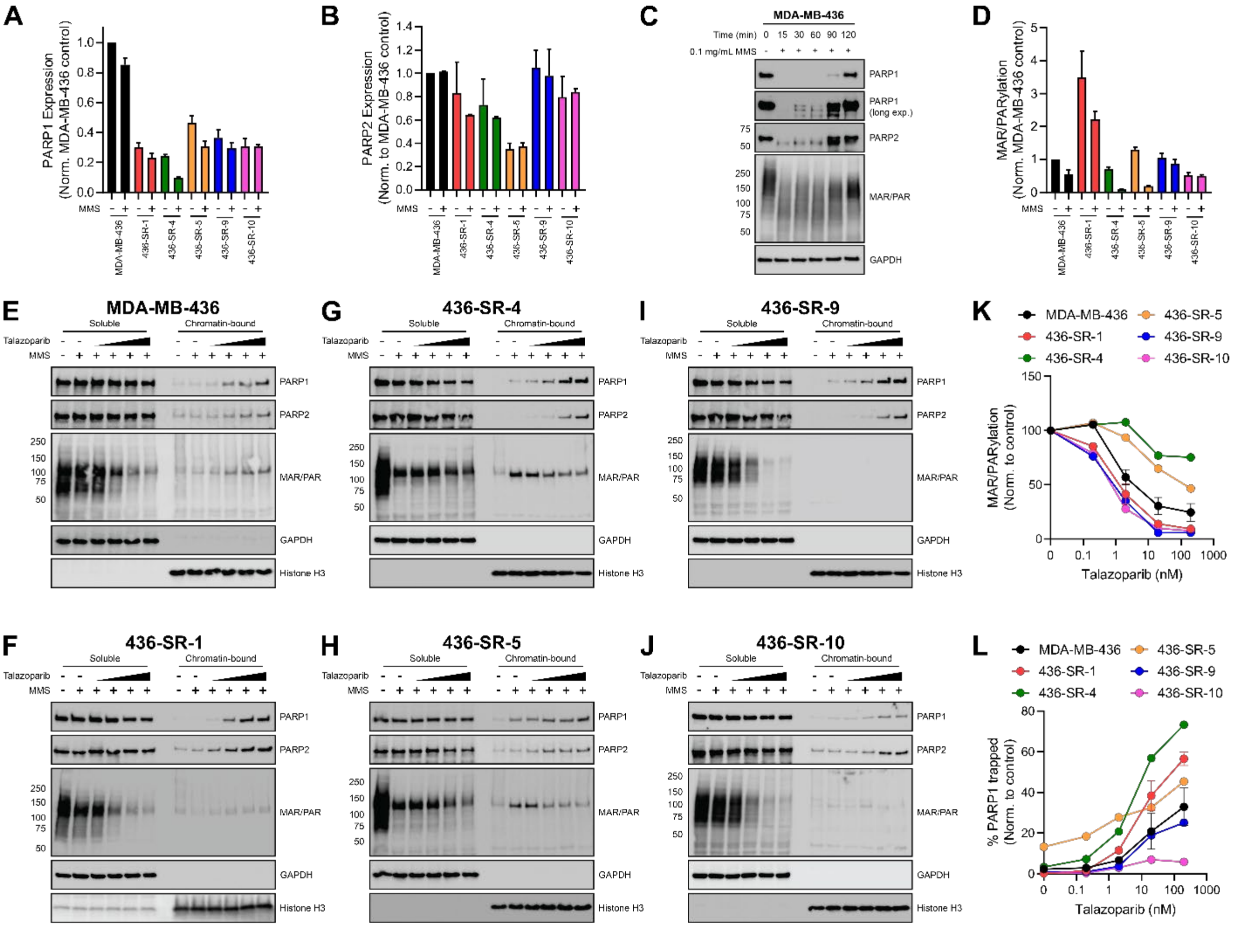


**Figure S4.** Quantification of **A)** PARP1 expression and **B)** PARP2 expression in MDA-MB-436 and SR cells from Figure 5A depicted as mean ± range from duplicate experiments. **C)** Western blot of soluble fractions from MDA-MB-436 cells treated with MMS over time. **D)** Quantification of MAR/PARylation in MDA-MB-436 and SR cells from Figure 5A depicted as mean ± range from duplicate experiments. Representative Western blots of soluble and chromatin-bound fractions from **E)** MDA-MB-436, **F)** 436-SR-1, **G)** 436-SR-4, **H)** 436-SR-5, **I)** 436-SR-9, and **J)** 436-SR-10 cells treated with DMSO control, 0.1 mg/mL MMS, and/or 0.2, 2, 20, 200 nM talazoparib for 2 hours. **K)** Quantification of talazoparib inhibition of MAR/PARylation in MDA-MB-436 and SR cells from panels C-H. **L)** Quantification of talazoparib-induced PARP1 trapping in MDA-MB-436 and SR cells from panels C-H.
